# NATF (Native And Tissue-specific Fluorescence): A strategy for bright, tissue-specific GFP labeling of native proteins

**DOI:** 10.1101/426205

**Authors:** Siwei He, Andrea Cuentas-Condori, David M. Miller

## Abstract

GFP labeling by genome editing can reveal the authentic location of a native protein but is frequently hampered by weak GFP signals and broad expression across a range cell types in multicellular animals. To overcome these problems, we engineered a <u>N</u>ative <u>A</u>nd <u>T</u>issue-specific Fluorescence (NATF) strategy which combines CRISPR/Cas-9 and split-GFP to yield bright, cell-specific protein labeling. We use CRISPR/Cas9 to insert a tandem array of seven copies of the GFP11 β-strand (*gfp11_x7_*) at the genomic locus of each target protein. The resultant *gfp11_x7_* knock-in strain is then crossed with separate reporter lines that express the complementing split-GFP fragment (*gfp1*-*10*) in specific cell types thus affording tissue-specific labeling of the target protein at its native level. We show that NATF reveals the otherwise undetectable intracellular location of the immunoglobulin protein, OIG-1, and demarcates a receptor auxiliary protein LEV-10 at cell-specific synaptic domains in the *C. elegans* nervous system.

## INTRODUCTION

Reliable localization of a given protein can provide useful clues to its mechanism of action. One way to achieve this goal is to label the protein of interest with tags, such as fluorescent proteins (e.g., GFP)^1,2^ or small peptides (e.g. FLAG, HA)^3^. Because tagged proteins are typically expressed with heterologous promoters or from multicopy transgenic arrays, this approach can result in misleading signals due to over-expression^4^. This problem can be obviated by using CRISPR-Cas9 for single copy labeling of the native protein^5,6^, but this genome editing strategy suffers from two additional limitations. First, the endogenous expression level of a target protein may be too low for detection. Second, the protein of interest may be expressed in several tissues thus preventing a clear delineation of cell-specific localization in multicellular organisms. Here we describe an experimental approach, NATF (**N**ative **A**nd **T**issue-specific **F**luorescence or “Native”) that exploits a combinatorial strategy to resolve both of these problems.

Our approach relies on the finding that the barrel-like GFP structure can be reconstituted by the spontaneous interaction of two separate GFP peptides derived from the highly stable GFP variant, superfolder GFP (sfGFP). The larger of these fragments is comprised of the first 10 β-strands (GFP1-10). Its smaller complement, a short, 16 amino acid sequence, contains the eleventh β-strand (GFP11). Neither GFP1-10 nor GFP11 fluoresce independently but a strong signal is restored in the hybrid split-GFP that they reconstitute^7^. Thus, to enhance the GFP signal, a target protein can be tagged with multiple copies of the short GFP11 peptide and then co-expressed with excess GFP1-10^8,9^. In addition to labeling the native protein with smaller covalent tags, this combinatorial approach offers the further benefit of limiting the GFP signal to the specific cell type in which GFP1-10 is expressed (**Figure 1a**).

**Figure 1.**
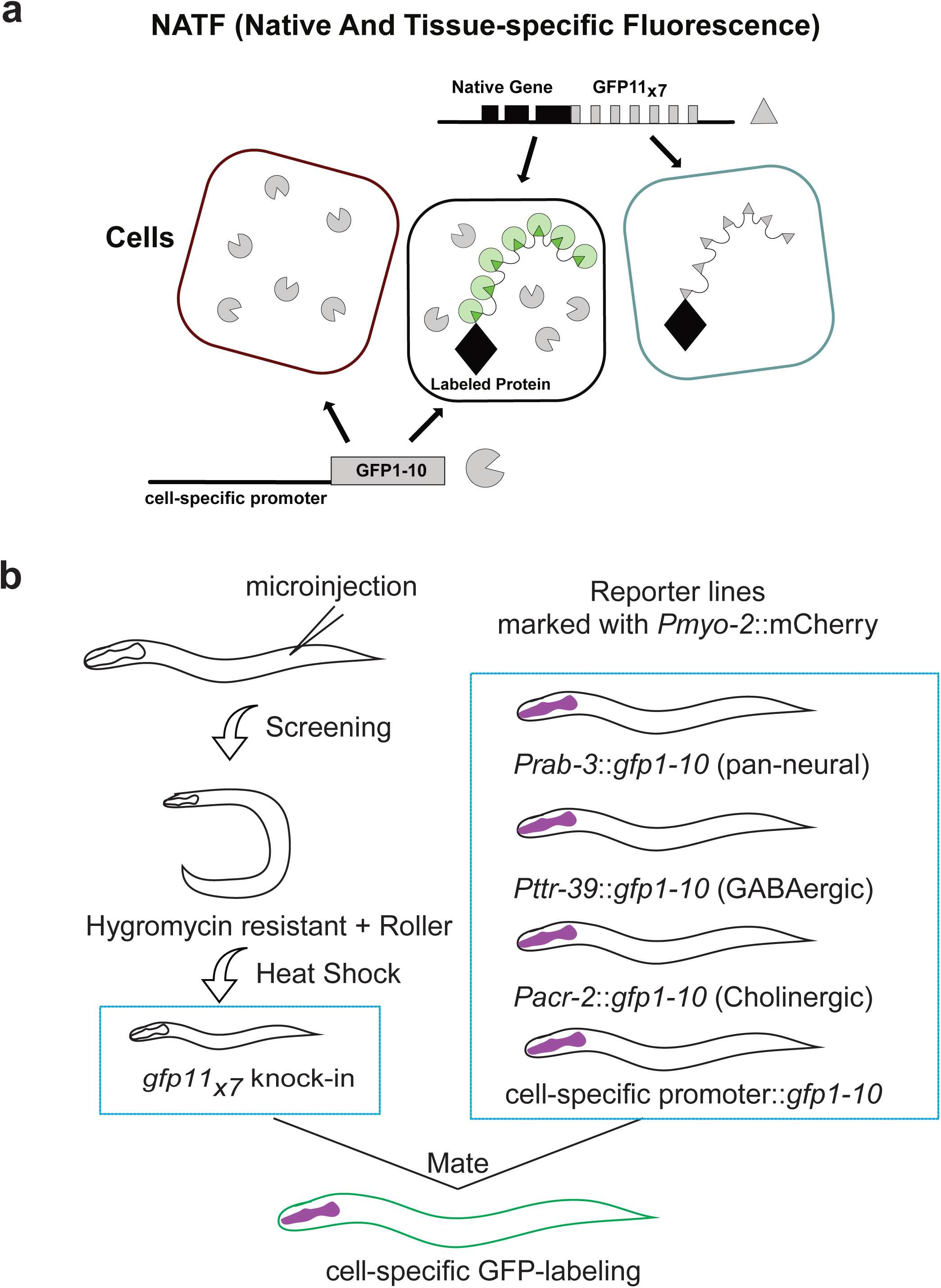
Robust, tissue-specific labeling of a target protein at its native expression level. **a.** NATF (Native And Tissue-specific Fluorescence). CRISPR/Cas-9-mediated gene editing is used to label a protein of interest with seven copies of GFP11 (GFP11_x7_). Transgenic expression of GFP1-10 with cell-specific promoters results in a bright, stable NATF fluorescent signal from multiple, reconstituted GFP molecules in specific tissues. **b.** NATF work flow. Worms are injected with sgRNA, repair template and co-injection markers. *gfp11_x7_* knock-in worms are recovered after heat shock-induced excision of positive-selection genes. Crossing the *gfp11_x7_* knock-in with GFP1-10 reporter lines results in tissue-specific labeling of the target protein with a reconstituted NATF-GFP tag.

In this report, we describe a NATF toolbox that combines split-GFP and CRISPR technology for live-cell imaging of labeled *C. elegans* proteins expressed at native levels. With this approach, a GFP11 multicopy DNA array (GFP11_X7_) is inserted into the target gene. The resultant knock-in line can then be crossed with separate reporter lines in which GFP1-10 is expressed in different cell types for tissue-specific visualization of the reconstituted NATF GFP (**Figure 1a**). We utilized this strategy to demonstrate effective enhancement of an otherwise weak signal from single copy fluorescent protein labeling of a key protein (OIG-1) in the *C. elegans* nervous system as well as the cell-specific resolution of a receptor accessory protein (LEV-10) at closely spaced but functionally distinct synapses.

## RESULTS

### Tool box and strategy for NATF GFP labeling

We used a previously described CRISPR/Cas-9 system for genome editing in *C. elegans*^10^. In this approach, homology arms flank a self-excising cassette that carries positive selection markers (*sqt*-*1*) for identifying transgenic worms (“rollers”) and drug resistance (hygR) for detecting CRISPR/Cas9-induced integrants^10^. A brief heat shock treatment induces excision of the marker cassette to restore wild-type movement (“non-roller”) (**Figure 1b**). For split-GFP experiments, we replaced the fluorescent protein sequence in the original repair template plasmid with a *gfp11_x7_* insert^8^. Homology arms of ~500bps were used for the two genes targeted (*oig*-*1* and *lev*-*10*) in this study (**Figure S1a**). We also constructed plasmids for expressing GFP1-10 in specific cell types including body muscles, all neurons, cholinergic neurons and GABAergic neurons (**Figure 1b** and **Figure S1b**). In these lines, the GFP1-10 transgenes are carried as extrachromosomal arrays that are maintained by selecting for pharyngeal coninjection marker (*Pmyo*-*2::*mCherry) (**Figure 1b**). Cell-specific drivers are flanked with multiple cloning sites to facilitate construction of plasmids for GFP1-10 expression in other tissues (**Figure S1b**). *gfp*11_x7_ knock-in strains can be confirmed within two weeks of the initial injection and then crossed with GFP1-10 expressing lines for characterization (**Figure 1b**).

### NATF GFP labeling reveals the intracellular localization of OIG-1 in GABAergic motor neurons

*oig*-*1* encodes a soluble protein with a single immunoglobulin domain (**Figure 2b**) that is temporally regulated in GABAergic motor neurons to antagonize a synaptic remodeling program; in *oig*-*1* mutants, a postsynaptic acetylcholine receptor (AChR) containing the AChR subunit, ACR-12::GFP, is ectopically relocated from dorsal to ventral GABAergic neuron processes. OIG-1 is secreted when over-expressed from multicopy transgenic arrays to produce bright puncta adjacent to clusters of ACR-12::GFP (**Figure 2a,b**)^11,12^. To ask if OIG-1 is also secreted when expressed from the native locus, we used CRISPR/Cas-9 to engineer a single copy knock-in of the red fluorescent protein, TagRFP together with a 3XFLAG epitope tag (**Figures 2b, S2a-c**). We used immunoblotting to confirm expression of TagRFP::3XFLAG::OIG-1 (**Figure 2c**) but failed to detect TagRFP fluorescence *in vivo* (**Figure 2d-g**). To produce a potentially brighter signal, we created a *gfp11_x7_::oig*-*1* knock-in (**Figure 2b**) with a sgRNA that targeted the same 5’-N_18_GGNGG site used for the TagRFP insert^13^. Successful knock-in of *gfp11_x7_* was confirmed by sequencing (data not shown). The resultant GFP11_X7_::OIG-1 fusion protein is likely functional as ACR-12::GFP puncta show robust localization to GABA neuron processes in both dorsal and ventral nerve cords as observed in wild type (**Figure S2 b-d**). We then crossed the *gfp11_x7_::oig*-*1* knock-in with a pan-neural *Prab*-*3::gfp1*-*10* transgenic line. Consistent with our previous findings, the OIG-1 signal can be detected in head neurons and in both dorsal and ventral nerve cords (**Figure 2h-k**)^11^. Co-localization of the OIG-1 NATF GFP signal with the pan-neural marker *Prab*-*3::*NLS::mCherry confirmed OIG-1 expression in neurons (**Figure 2l**). As an independent strategy to validate OIG-1 expression in GABA neurons, we crossed the *gfp11_x7_::oig*-*1* line with *Pttr*-*39::gfp1*-*10* which is selectively expressed in DD and VD class GABAergic motor neurons^14^. In this case, the OIG-1 NATF GFP signal is limited to VD neurons with either weak or undetectable expression in DD neurons at the L4 larval stage (**Figure 2l**). This finding confirms previous results obtained with a *Poig*-*1::gfp* transcriptional reporter that was expressed in VD but not DD neurons after the L2 larval stage^11^. Because the GFP1-10 peptide is expressed intracellularly in these strains, the NATF GFP signal likely derives from cytoplasmic OIG-1. Notably, the OIG-1 NATF GFP signal is diffusely visible throughout VD neuron soma and neurites (**Figure 2h-m**) and does not show the distinctive punctate appearance of over-expressed OIG-1 from a multicopy array (**Figure 2a, b**). In addition, we failed to detect extracellular NATF GFP when the *gfp11_x7_::oig*-*1* knock-in was crossed with a transgenic line in which the GFP1-10 peptide is secreted from neurons (*Prab*-*3::ss::gfp1*-*10*) **(Figure S3a)**. As a positive control, we showed that the secreted form of GFP1-10 in the *Prab*-*3::ss::gfp1*-*10* strain is functional because it robustly labels a GFP11 peptide fused to the extracellular domain of the synaptic membrane protein, NLG-1^15^ (**Figure S3b-d**). Thus, we conclude that OIG-1 is not secreted when expressed at the native level but localizes intracellularly (manuscript in preparation). This surprising finding critically depended on the use of the NATF strategy to detect low levels of native OIG-1 expression and thereby circumvent artifactual localization due to OIG-1 over-expression from multicopy arrays.

**Figure 2.**
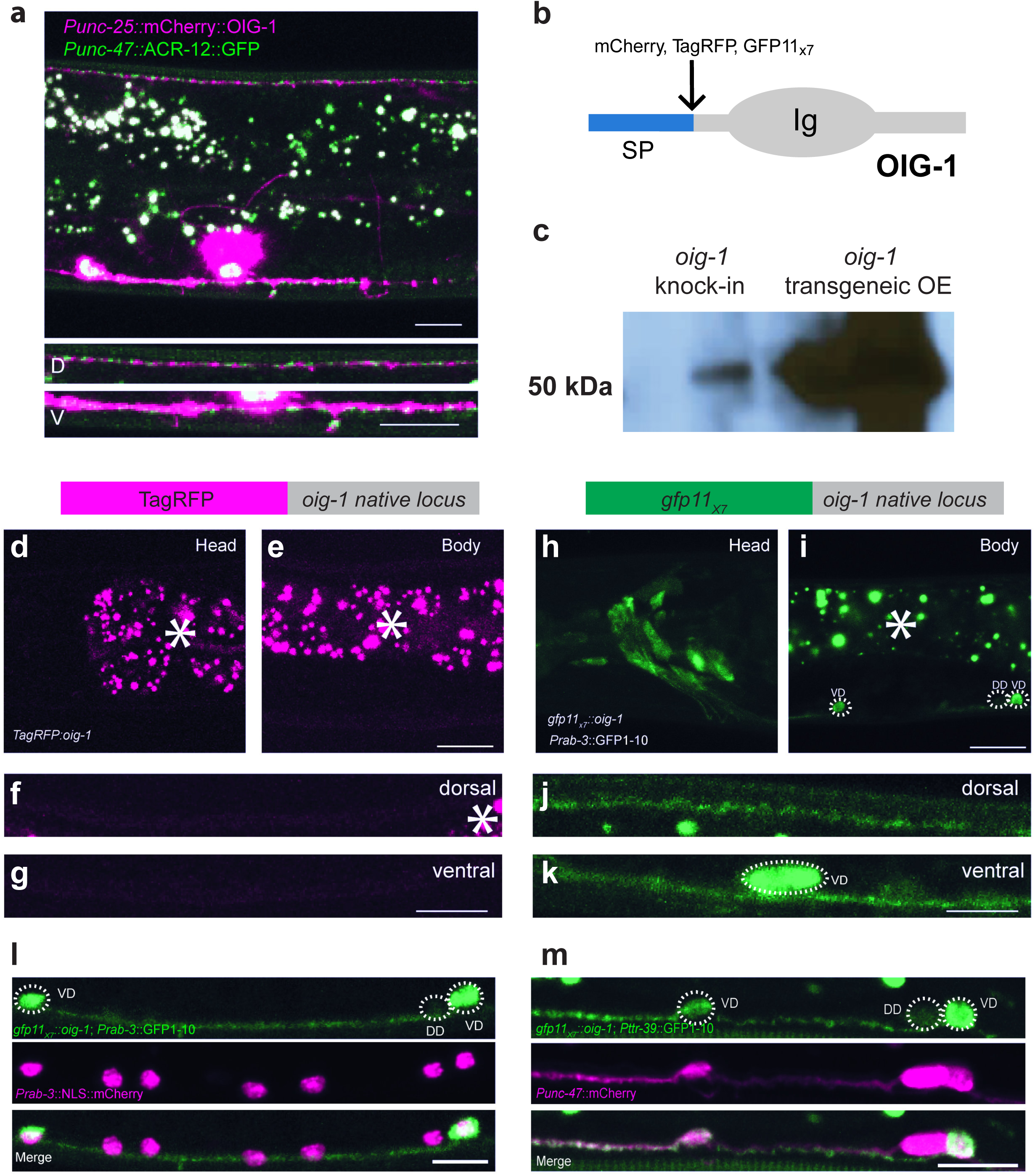
CRISPR knock-in of GFP11_x7_ in OIG-1 reveals its intracellular localization in GABAergic motor neurons. **a.** Over-expression of mCherry::OIG-1 from a transgenic array (*Punc*-*25::mCherry::oig*-*1*) produces bright puncta adjacent to ACR-12::GFβ-labeled postsynaptic clusters of AChRs in GABAergic motor neurons (*Punc*-*47::acr*-*12::gfp*). Insets show dorsal (D) and ventral (V) nerve cords. Scale bars are 10μm. **b.** OIG-1 protein showing N-terminal signal peptide (SP), C-terminal Immunoglobulin domain (Ig) and insertion site for fluorescent tags (mCherry, TagRFP, GFP11_x7_). **c.** Immunoblot stained for 3XFLAG tag detects expression of single copy TagRFP::OIG-1 knock-in and confirms over-expression (OE) of mCherry::3XFLAG::OIG-1 from multicopy transgenic array. **d-g.** TagRFP knock-in at the *oig*-*1* locus (top) does not result in detectable TagRFP::OIG-1 fluorescence. Note absence of TagRFP::OIG-1 signal in head region **(d),** body **(e)** and dorsal **(f)** and ventral **(g)** nerve cords. L4 larvae. Asterisks mark autofluorescent granules. **h-k.** OIG-1 expression in the nervous system. The *gfp11_x7_::oig*-*1* knock-in (top) was crossed with the pan-neural transgenic line expressing *Prab*-*3*::GFP1-10. A diffuse OIG-1 NATF-GFP signal is detected in a head neurons **(h)** and in VD but not DD GABAergic motor neurons in the ventral cord **(i)**. NATF-GFβ-labeled OIG-1 is detectable in both dorsal and ventral nerve cords **(j, k)**. Asterisk marks autofluorescent granules. Scale bars for **d-k** are 10 μm. **l-m.** OIG-1 expression in the nervous system and in GABAergic motor neurons. The *gfp11_x7_::oig*-*1* knock-in line was crossed with the pan-neural marker *Prab*-*3::gfp1*-*10* and all neurons labeled with a nuclear-localized pan-neural marker *Prab*-*3*::NLS::mCherry **(l)**. The *gfp11_x7_::oig*-*1* knock-in was crossed with the GABAergic motor neuron-specific marker *Pttr*-*39::gfp1*-*10* and all GABA neurons were labeled with *Punc*-*47*::mCherry **(m)**. Note the OIG-1 NATF-GFP signal in VD but not DD motor neurons^11^. GFP1-10 is cytosolically expressed from the *Prab*-*3::gfp1*-*10* and *Pttr*-*39::gfp1*-*10* transgenes thus indicating OIG-1 is intracellularly localized at its native expression level. All images were obtained from L4 stage larvae. Scale bars are 10 μm.

### NATF GFP labeling reveals discrete locations for the CUB domain protein LEV-10 in different cell types

Having shown that NATF could detect a soluble protein (OIG-1), we next targeted, LEV-10, a CUB domain transmembrane protein that clusters AChRs at postsynaptic sites in body muscles^16^. First, we created a CRISPR/Cas9 knock-in line in which a single copy of GFP was fused to the intracellular C-terminus of LEV-10 (see **Figure 4a**). We detected LEV-10::GFP in both ventral and dorsal nerve cords as predicted for a protein that localizes to body muscle synapses^16^. LEV-10::GFP puncta were also detected in the head region where motor neurons synapse with body muscles on the inside surface of the nerve ring^17,18^ (**Figure 3a**). For NATF GFP labeling of body muscle synapses, we generated a *lev*-*10::gfp11_x7_* knock-in and crossed it with a muscle-specific transgenic line expressing GFP1-10 (*Pmyo*-*3::gfp1*-*10*). The LEV-10 NATF GFP signal in the head region and axial nerve cords (**Figure 3b**) mimics that of the single copy *lev*-*10::gfp* knock-in but is noticeably brighter (**Figure 3a**). We quantified the GFP signal for each marker at the nerve ring muscle synapses to confirm that the LEV-10 NATF fluorescence is ~3X brighter than the GFP signal from the *lev*-*10::gfp* single copy insertion^8^ (**Figure 3c**). In addition to determining that the *lev*-*10::gfp11_x7_* array yields a stronger signal than that of the single copy *lev*-*10*::GFP insert, we also showed that NATF GFP is more resistant to photobleaching as previously demonstrated for reconstituted split-GFP from measurements *in vitro*^8^ (**Figure 3d**).

**Figure 3.**
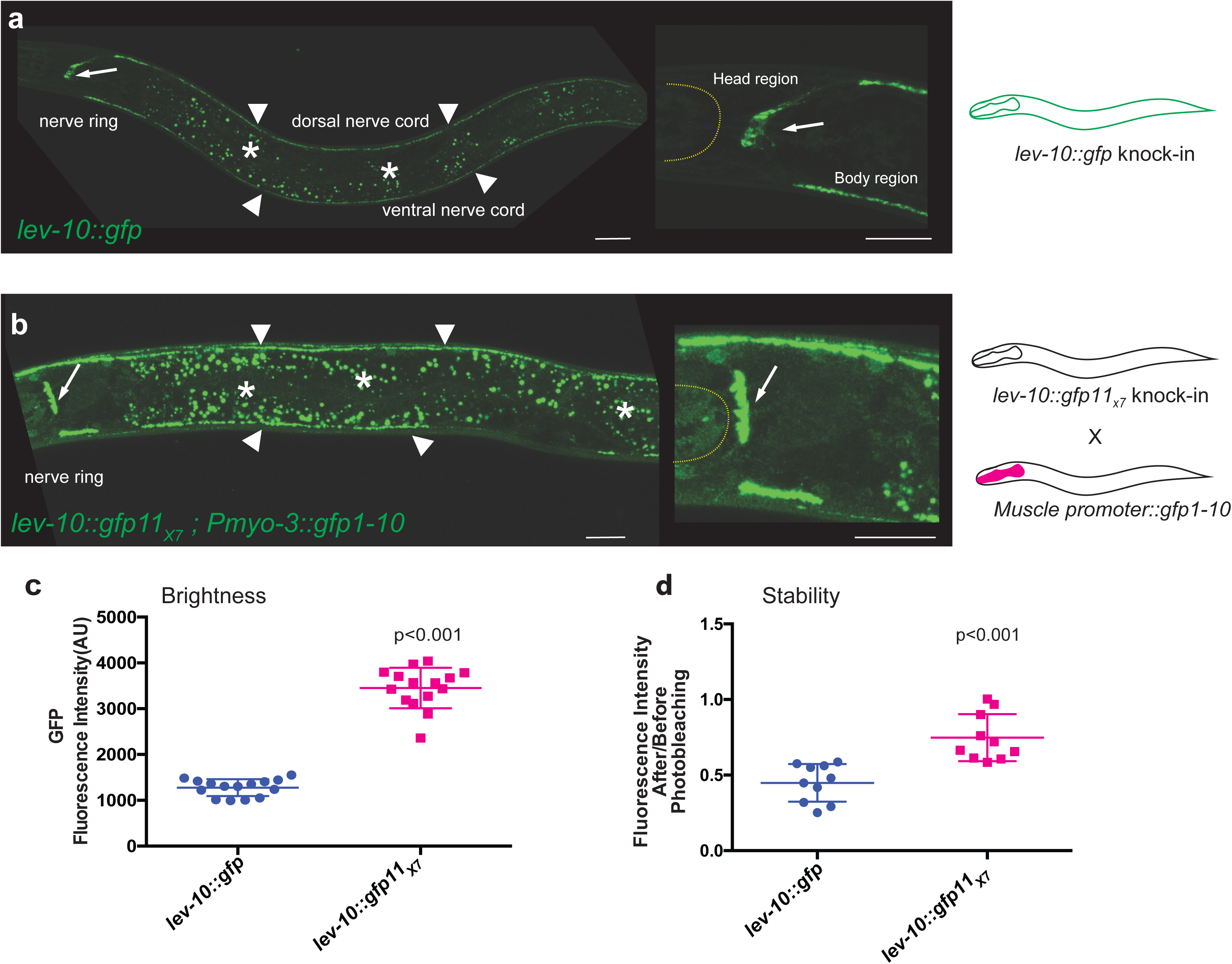
*Iev*-*10::gfp11_x7_* yields a stronger NATF GFP signal than the single copy *lev*-*10::gfp* knock-in at synapses in neurons and muscle cells. **a.** Confocal Images showing localization of LEV-10::GFP in a single-copy GFP knock-in at the native *lev*-*10* gene (*lev*-*10::gfp*). LEV-10::GFP puncta are visible at the nerve ring (arrow) and in ventral and dorsal nerve cords (arrowheads) Asterisks mark gut autofluorescence. Inset (right) shows anterior region of image on left. **b.** Confocal images of the NATF GFP signal at body muscle synapses arising from the combination of the *lev*-*10::gfp11_x7_* knock-in with *Pmyo*-*3::gfp1*-*10*. Dashed line marks the anterior pharyngeal bulb. NATF GFP (arrow) is detected at neuromuscular synapses near the nerve ring. Scale bars are 20 μm. **c**. NATF GFP at body muscle synapses in the nerve ring labeled with *lev*-*10::gfp11_x7_* is significantly brighter (~3X) than the single copy *lev*-*10::gfp* knock-in. p < 0.001, N = 15, Student’s T-test. **d.** NATF GFP at body muscle synapses in the nerve ring labeled with *lev*-*10::gfp11_x7_* is significantly more stable to photobleaching than the single copy *lev*-*10::gfp* knock-in, N = 10, p < 0.001, Student’s T-test. See Methods.

**Figure 4.**
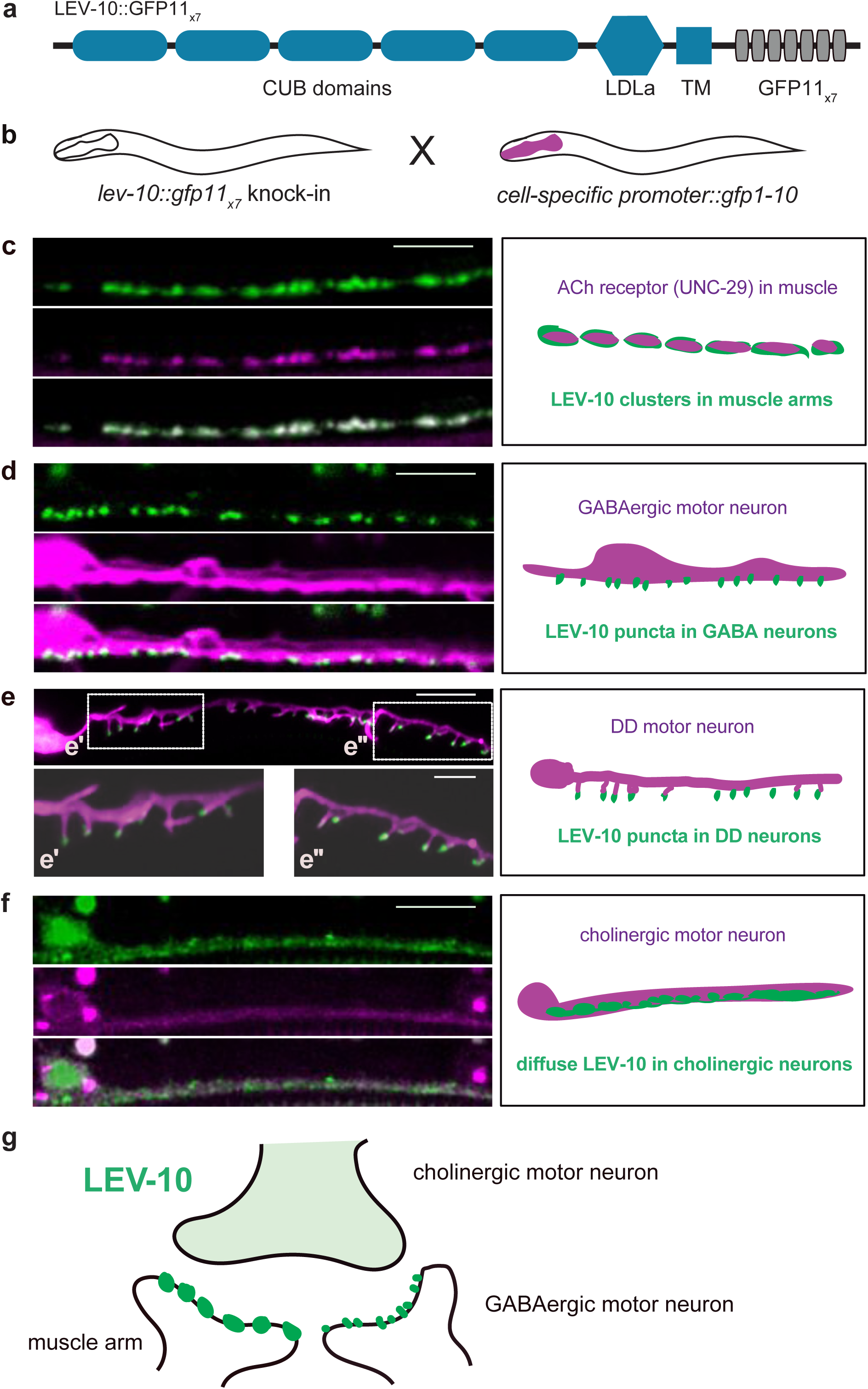
Visualization of NATF LEV-10 at cell-specific synapses. **a.** Schematic of LEV-10::GFP11_x7_ showing extracellular CUB and LDLa domains with transmembrane (TM) region and cytoplasmic tail with GFP_11×7_ insert. **b.** Cell-specific labeling strategy: The *lev*-*10:: gfp11_x7_* knock-in is crossed with separate transgenic lines expressing GFP1-10 in specific cell types. **c-f.** Representative images (left) and schematics (right) of ventral nerve cord region of L4 larvae showing NATF GFP arising from complementation of the *lev*-*10:: gfp11_x7_* knock-in with cell-specific expression of GFP1-10: **(c)** *Pmyo*-*3::gfp1*-*10* (body muscle), **(d)** *Pttr*-*39::gfp1*-*10* (DD and VD GABAergic motor neurons), **(e)** *Pflp*-*13::gfp1*-*10* (DD GABAergic motor neurons) Super-resolution images with insets. **(f)** *Pacr*-*2::gfp1*-*10* (cholinergic motor neurons). *Pmyo*-*3::unc*-*29:TagRFP* marks ACh receptors from muscle in **c**; *Punc*-*47*::mCherry labels GABA neurons in d; *Pflp*-*13::LifeAct::mCherry* marks DD neurons in **e** and *Punc*-*4::mCherry* labels cholinergic motor neurons in **f.** Scale bars are 5 μm **(c - f)** and 2 μm **(e’ and e”)**. **g.** Schematic showing distribution of LEV-10 NATF GFP at a dyadic synapse of presynaptic cholinergic motor neurons with postsynaptic body muscles and GABAergic motor neurons in the ventral nerve cord.

In addition to expression in muscle, our independent studies have shown that LEV-10 is also expressed in ventral cord neurons where it co-localizes with AChRs at postsynaptic sites in GABAergic motor neurons (manuscript in preparation). In the motor neuron circuit, cholinergic motor neurons form dyadic synapses that innervate closely spaced postsynaptic domains in body muscle and GABA neurons (see **Figure 4g**)^17^. Both of these postsynaptic regions in the ventral nerve cord region should be labeled in the *lev*-*10::gfp* knock-in and thus, cannot be unambiguously identified (**Figure 3a**). To resolve this problem, we crossed the *lev*-*10::gfp11_x7_* knock-in with transgenic lines that express GFP1-10 in either body muscles (*Pmyo*-*3::gfp1*-*10*) or in DD and VD GABAergic motor neurons (*Pttr*-*39::gfp1*-*10*). NATF GFP puncta can be readily detected in both cases (**Figures 4c,d**) but are brighter in muscles than in GABAergic neurons (data not shown). Expression of a TagRFP-labeled AChR subunit, UNC-29, in muscle confirms co-localization of UNC-29::TagRFP and LEV-10 NATF GFP (**Figure 4c**). Expression of GFP1-10 in DD and VD neurons produces LEV-10 NATF GFP puncta that overlap with a cytoplasmic GABA neuron mCherry marker (*Punc*-*47::*mCherry) as predicted for LEV-10 protein that localizes to GABA neuron synapses (**Figure 4d**). To confirm the postsynaptic location of LEV-10 in GABA neurons, we used a DD-specific construct (*Pflp*-*13::gfp1*-*10*) to generate a LEV-10 NATF GFP signal. In this case, super-resolution imaging resolves distinct LEV-10 NATF GFP puncta at the tips of postsynaptic spine-like projections that have been recently described in the ventral processes of mature DD neurons (**Figure 4e and Figure S4**)^19^. Notably, we have also observed that the AChR marker, ACR-12::GFP, is positioned in the same distal location in DD dendritic spines and that these spines are aligned with presynaptic cholinergic vesicles (manuscript in preparation)^19^. In addition to resolving LEV-10 localization at distinct postsynaptic locations in muscle *vs* GABA neurons, we also used a cholinergic motor neuron driver (*Pacr*-*2::gfp1*-*10*) to detect a separate LEV-10 NATF signal in ventral cord cholinergic neurons. In this case, LEV-10 NATF GFP is diffuse (**Figure 4f**) and asymmetrically localized to the ventral but not dorsal nerve cord (data not shown), a labeling pattern that closely resembles the perisynaptic position of the AChR subunit ACR-12::GFP in cholinergic motor neurons^20^. Because LEV-10 is expressed at its native level and retains its AChR clustering function (data not shown) when fused to the GFP11_X7_ adduct, it seems likely that each of the three distinct, cell-specific LEV-10 NATF signals that we have detected with our combinatorial approach (i.e., muscle, GABA neurons, cholinergic neurons) marks authentic subcellular locations for the endogenously expressed LEV-10 protein.

## DISCUSSION

Although our results have determined that fusion with the GFP11_X7_ domain does not disrupt either OIG-1 or LEV-10 activity, other proteins may be less tolerant. In that event, smaller adducts with fewer copies of GFP11 could be attempted^21^. In that case, GFP signal augmentation will be diminished but tissue-specific labeling is still possible. Because the gfp11_×7_ insert is stably integrated at the native locus and is thus limiting, the complementing GFP1-10 peptide can be provided from multicopy transgenic arrays without risk of inducing over-expression artifacts. Thus, a given GFP*11_X7_* split GFP insert can be rapidly tested with multiple tissue specific GFP1-10 transgenic lines which can be readily generated using conventional methods. A similar combinatorial approach should also be useful for tissue-specific protein labeling in other model organisms. We note that NATF can be modified for multicolor split-GFP imaging with cyan (CFP) and yellow (YFP) GFP variants or with the sfmCherry marker^11,12^.

## ACKNOWLEDGEMENTS

We thank members of the Miller lab for critical reading of the manuscript, Sierra Palumbos and Alice Siqi Chen for plasmid construction, Lakshmi Sundararajan for help with confocal imaging and Oliver Hobert for sharing strains. Some nematode strains used in this work were provided by the Caenorhabditis Genetics Center, which is funded by the NIH National Center for Research Resources (NCRR). Super-resolution imaging was acquired at the Vanderbilt Cell Imaging Shared Resource (1S10OD201630-01). This work was supported by National Institutes of Health grants to DMM (R01NS081259 and R01NS106951). ACC is supported by an AHA predoctoral fellowship (18PRE33960581).

## AUTHOR CONTRIBUTIONS

SH, ACC and DMM conceived the project. SH and ACC performed the experiments and analyzed the data. SH, ACC and DMM wrote the manuscript.

## MATERIALS AND CORRESPONDENCE

David M. Miller, Ph.D

## Methods

### *C. elegans* strains

*C. elegans* strains were maintained at room temperature on NGM plates seeded with OP50 as previously described^22^. Some strains were obtained from the *Caenorhabditis* Genetics Center (CGC). The N2 Bristol strain was used as the wild-type reference. Transgenic animals were generated using standard microinjection techniques. Unless noted otherwise, 100 ng/μL total DNA injection samples were prepared with the pBluescript plasmid, as carrier. The strains used in this study are described in Supplementary Table 1.

### Molecular biology

#### sgRNA/Cas-9 plasmid design

A 200bp DNA sequence that contains the desired cut site was submitted to opitimized CRISPR Design online tool (http://crispr.mit.edu/) to predict sgRNA sequences. To enhance gene editing efficiency, we selected a 5’ N_18_GGNGG sequence^13^ as a sgRNA targeting site for both *oig*-*1* and *lev*-*10*. For *oig*-*1*, GGAGAGAAAGACGAAAATGG was cloned into pDD162(Addgene #47549), a plasmid that contains the sgRNA backbone and Cas-9 expression system using Q5 site directed mutagenesis (NEB). Similarly, for *lev*-*10*, ACGAATCGACTGGTGGCCGG was used as the sgRNA binding sequence, which is ~80bp upstream of the *lev*-*10* stop codon.

#### CRISPR repair template for *oig*-*1* and *lev*-*10*

To create the SEC repair template for *oig*-*1* TagRFP CRISPR knock-in, flanking ~500bp genomic DNA regions immediately upstream and downstream of the desired insertion site were amplified by PCR using the following primers (Primer 1 and Primer 2 for upstream homology arm and Primer 3 and Primer 4 for downstream homology arm) with overlap regions to the target plasmid pDD284(Addgene #66825):

OIG-1 Primer 1: 5’ gacgttgtaaaacgacggccagtcgacctaaccattccaaaagat

OIG-1 Primer 2: 5’ tgagctcctctcccttggagaccatcgcatttattccaactgata

OIG-1 Primer 3: 5’ ttacaaggatgacgatgacaagagaaaatcttcgcatatagaaga

OIG-1 Primer 4: 5’ caggaaacagctatgaccatgttatccaagtcggagtactgttca

The amplified DNA fragments were cloned into plasmid pDD284 using Gibson cloning (NEB) to create the repair template. The corresponding PAM sequence in the repair template plasmid was mutated from AGG to CCC using Q5 site-directed mutagenesis to produce the final plasmid, pSH30. Correct insertions and mutations were confirmed by sequencing.

To create the SEC repair template (pSH55) for the *oig*-*1* GFP11_x7_ CRISPR knock-in, the GFP11_x7_ coding sequence was amplified from a previously published plasmid (Addgene #60910) and inserted into pSH30 to replace the TagRFP sequence by In-Fusion cloning (Takara) with the following primers.

Fragment.FOR 221 5’ gttggaataaatgcgatgcgtgaccacatggtcctt

Fragment.REV 222 5’ aaagtacagattctcggtgataccggcagcat

Vector.FOR 223 5’ gagaatctgtactttcaatccggaaaggtaag

Vector.REV 224 5’ cgcatttattccaactgatagaaagcataaaagtagt

To create the GFP and GFP11_x7_ knock-in repair template for LEV-10, we designed a two-step In-Fusion cloning method. ~500bp of DNA sequences upstream and downstream of the *lev*-*10* stop codon were selected for flanking homology arms. DNA was amplified and then sequentially cloned into pSH30 or pSH55 to replace the original *oig*-*1* homology arms. The resultant plasmids were then used as templates for site-direct mutagenesis to create the final repair template plasmid with sgRNA binding sequences mutated, pSH84(GFP knock-in) and pSH85(GFP11_x7_ knock-in). The primers for these cloning steps were designed using the same strategy as described above. Primer sequences are available on request.

#### GFP1-10 reporter plasmids

The DNA sequence of GFP1-10 was amplified from pcDNA3.1-GFP1-10 (Addgene #70219) and cloned into pGH8 (*Prab*-*3*:: mCherry) using infusion cloning to create pSP1(*Prab*-*3::gfp1*-*10*). The DD and VD GABAergic neuron-specific promoter, *Pttr*-*39*, the cholinergic specific promoter, *Pacr*-2, and muscle-specific *Pmyo*-*3* promoter were amplified to replace the *Prab*-*3* promoter in pSP1 to create pSH79(*Pttr*-*39::gfp1*-*10*), pSH88(*Pacr*-*2::gfp1*-*10*), pSH86(*Pmyo*-*3::gfp1*-*10*) and pSH87(*Pflp*-*13::gfp1*-*10*). To create a secreted GFP1-10 construct, infusion cloning was used to add the first 114bp of the *oig*-*1* sequence including the signal sequence^11^ prior to the start codon of GFP1-10 in pSP1. The combined sequence was analyzed using SignalP 4.1 Server to confirm that the predicted signal peptide was intact. The final plasmid, pSH69 (*Prab*-*3*::ssGFP1-10), was confirmed by sequencing.

### Confocal microscopy and image processing

Images of fluorescently-labeled worms were captured at room temperature in live *C. elegans* using a Nikon A1R confocal microscope. Nematodes were immobilized with 15mM levamisole/0.05% tricaine on a 2% agarose pad in M9 buffer. All images for ACR-12::GFP fluorescence quantification were obtained with the same settings using the 40X oil, NA 1.3 objective and Nyquist collection. Constant laser power was used to compare the LEV-10::GFP fluorescence intensity to that of the NATF GPF signal produced by the combination of LEV-10::GFP11_x7_ with *Pmyo*-*3*::GFP1-10. Images in **Figure 4** were 3D-deconvolved with NIS-Elements with Automatic algorithm. For other images, ND2 files generated with NIS-Elements were imported into Fiji for analysis. Maximum intensity projections were generated by selecting stacks that have both ventral and dorsal signals. The mean fluorescence intensity of each animal after subtracting background was used for statistical analysis. To measure the stability of GFP signal in *lev*-*10::gfp* and *lev*-*10::gfp11_x7_*; *Pmyo*-*3*::GFP1-10 lines, a region of interest (ROI) of the same size was bleached with a 405nm laser for 15 seconds at 50% laser power. Images of the ROI were collected and compared before and after photo bleaching.

### Airy Scan Imaging

Worms were mounted on 10% agarose pads and immobilized with 15mM levamisole/0.05% tricaine dissolved in M9. A Zeiss LSM880 microscope equipped with an AiryScan detector and a 63X/1.40 Plan-Apochromat oil objective lens was used to acquire super resolution images of the DD neuron (Figure 4e). Images were acquired as a Z-stack (0.19μm/step), spanning the total volume of DD neuron and submitted for AiryScan image processing using ZEN software.

### Statistical Analysis

For all experiments, sample numbers were n >10. Student’s t-test was used for comparison between two groups. P < 0.05 was considered significant. Prism 6 was used for statistical analysis.

